# Response of mycorrhizae to herbivory and soil moisture in a semiarid grazing ecosystem

**DOI:** 10.1101/2020.08.13.248237

**Authors:** Yadugiri V Tiruvaimozhi, Sumanta Bagchi, Mahesh Sankaran

## Abstract

Arbuscular mycorrhizal fungal (AMF) symbioses with plants can be influenced by top-down forces such as grazing, and also by bottom-up forces such as soil resource availability, both of which are being altered by anthropogenic and global change drivers. While the influence of each of these factors on AMF symbioses has been widely studied, explicit tests of the relative strengths of top-down versus bottom-up influences on these ubiquitous plant root symbioses are few. We studied AMF colonization responses of four species of graminoids (3 grasses *Elymus longae-aristatus*, *Leymus secalinus* and *Stipa orientalis*, and a sedge *Carex melanantha*) common to semiarid high-altitude rangelands of the Spiti region, Trans-Himalaya, to changes in a top-down driver, grazing intensity (through short-term clipping and long-term grazer exclusion experiments), and a bottom-up driver, water availability (using irrigation treatments, and by evaluating responses to annual precipitation levels across years). Over three years, AMF colonization in all four host species was influenced by precipitation, with the highest and lowest AMF colonization levels corresponding to years with the lowest and highest rainfall, respectively. However, responses to long-term grazer exclusion differed among host species, and across years: while some species showed decreases in AMF colonization levels under grazing, others showed increases from ungrazed control levels, and these responses changed, even reversed, across years. Responses to short-term clipping and irrigation treatments also differed among hosts, with some species responding to irrigation alone, some to clipping and irrigation combined, and others showing no changes in AMF colonization from control levels in any of the treatments. In our study, long-term changes in water availability influenced AMF colonization levels, while short-term responses were host specific. Responses to above-ground tissue loss, however, differed among host species both in the long- and short-term. Overall, this study demonstrates that while AMF colonization levels correspond to annual precipitation levels in this semiarid ecosystem, host species also play a role in influencing plant-AMF interactions in these rangelands, with colonization levels and responses to abiotic factors changing with host species.

## Introduction

Arbuscular mycorrhizal fungi (AMF) play a major role in the cycling and storage of carbon (Treseder & Allen 2000) and the movement of nutrients and other resources through ecosystems. About 70-90% of all land plants host AMF in their roots for better access to nitrogen (N), phosphorus (P), water and other resources (Parniske 2008). In return, plants may invest 5-85% of their photosynthetically fixed carbon (C) in these fungi (Treseder & Allen 2000), with several factors affecting the level of investment. Investment in AMF can differ across host species, depending on the host plant’s position in the facultative-obligate mycorrhizal continuum (Moora 2014), or across ecosystems, with grasslands, in general, having the highest AMF root colonization levels (Treseder & Cross 2006). Investment in AMF can also change with time, with AMF colonization levels affected by factors such as changes in resource availability due to disturbances like grazing (Barto & Rillig 2010) and other global change drivers (Mohan et al. 2014).

AMF colonization levels can be influenced by a number of factors that affect host plant C fixation and retention. These include top-down forces such as herbivory and bottom-up forces such as soil resource availability (the usage of the terms ‘top-down’ and ‘bottom-up’ forces is *sensu* Wardle et al. 2004; Bardgett & Wardle 2010, where top-down refers to grazer (or other plant consumer)-mediated control and bottom-up to soil resource availability-mediated control on plant C allocation strategies). Defoliation due to herbivory can deplete plant C reserves and lead to decreased investment in AMF resulting in lower mycorrhizal colonization levels (Allsopp 1998; Barto & Rillig 2010). Alternatively, grazing can lead to higher nutrient and water demand for regrowth of host plant tissue, and result in increased plant allocation of C to AMF, and increased colonization levels (Antoninka et al. 2015). Bottom-up forces, such as increased nutrient and water availability, are expected to reduce plant requirement for extensive resource acquisition machinery such as AMF, and lead to lower colonization levels (Allen et al. 2003; Johnson 2010). The relative importance of top-down versus bottom-up forces in influencing AMF colonization, however, is not well understood.

What happens to AMF colonization when both herbivores and resource availability strongly shape vegetation growth, as in several grazing ecosystems that occur in dry regions? Here, we develop and test a predictive framework for levels of AMF colonization of plant roots under such conditions based on the functional equilibrium model (Brouwer 1963; Johnson et al. 2006). The functional equilibrium model adopts a plant perspective to predict plant C allocation strategies, even to symbionts such as AMF. This is reasonable because, while it is true that both the plant and fungal partners exert an influence in plant-AMF associations (Kiers et al. 2011; Fellbaum et al. 2012), the plant is still the more ‘dominant’ partner. Mycorrhizae are obligate biotrophs and have zero fitness in the absence of a plant partner, while plants have other mechanisms, such as roots, to access soil resources and can survive for some time without AMF (Johnson 2010). The underlying premise of the functional equilibrium model is that plant investment in C fixation and soil resource acquisition compete for the same internal resources in plants. Plant will invest in C acquisition (leaves) versus soil resource assimilation machinery (roots and AMF) to reduce the imbalance between C versus soil resource demand. In other words, this model suggests that plants should invest most in structures that improve access to the most limiting resource (Brouwer 1963; Bloom et al. 1985; Wilson 1988; Johnson 2010). When resources are limiting but grazing pressure is low (Fig. 1, Region I), plants might be expected to invest in AMF, i.e, have high AMF colonization levels, to better access soil resources, provided light is not limiting. Similarly, when resources are low, but grazing pressure is high (Fig. 1, Region II), AMF colonization levels can be expected to be high due to increased nutrient demand to support plant regrowth following herbivory. On the other hand, when resources are plentiful, plant investment in AMF is expected to be low, irrespective of herbivory levels (Fig. 1, Regions III and IV), since plants would allocate C aboveground rather than to nutrient acquisition machinery like AMF.

**Fig. 1.**
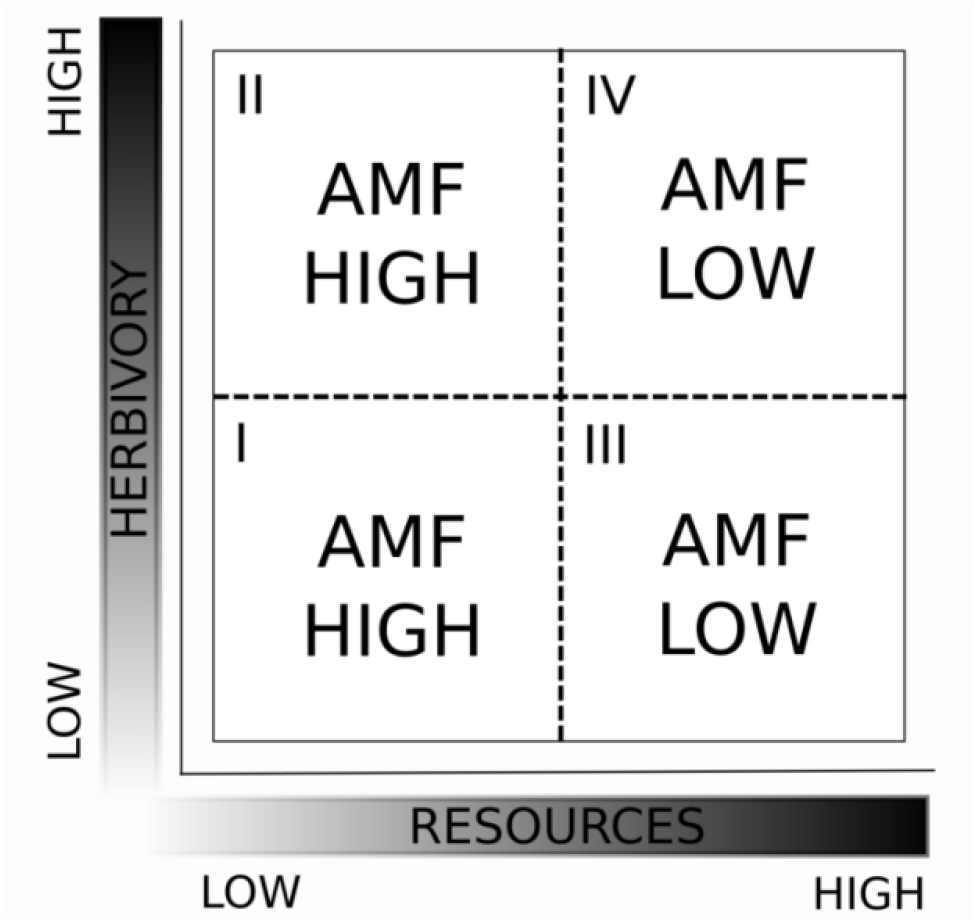
Predictive framework for the influence of top-down versus bottom-up forces on AMF colonization under low and high resource availability and herbivory pressure. The x-axis represents a gradient of resource availability going from low to high (light grey to dark grey), and the y-axis represents a similar gradient of herbivory pressure. The Roman numerals (I-IV) denote ‘Regions’ representing four possible combinations of resource availability and herbivory pressure.

The predictive framework described above is robust over temporal (e.g. short-term vs. long-term grazing) and spatial scales (e.g. high precipitation, where light might also be a limiting factor, vs. low precipitation sites, where light is not limiting). For instance, under low soil resource conditions, both chronically grazed and short-term experimentally clipped plants can be expected to invest C in belowground resource acquisition machinery such as AMF, to get enough water and nutrients to be able to replace grazed tissue. Under high resource conditions, the functional equilibrium model predicts that both these plants will invest C aboveground to compensate grazed tissue, while investment to belowground resource acquisition machinery will be minimal. Similarly, in high precipitation regions where light is limiting, vs. low precipitation regions, the predictions in Fig. 1 will hold, provided soil nutrients are not limiting. In the high precipitation site, in the absence of grazing, light is the most limiting resource. The functional equilibrium model would suggest that plant C investment in this case would be in aboveground tissues (for C capture), and AMF colonization would be low. In the presence of grazing, light limitation might be reduced, but since water is not limiting, plant C investment would be towards regrowing aboveground grazed tissue, and AMF colonization would still be low. In the low precipitation site, light is not limiting, and the scenarios we have described earlier would hold as is.

Several studies report effects of either herbivory (e.g. Barto & Rillig 2010, and studies referred therein) or altered resource availability (e.g. Augé 2001; Treseder 2004; Mohan et al. 2014, and studies referred therein) on AMF colonization, and these generally report much heterogeneity in responses. However, explicit tests of top-down versus bottom-up controls on AMF colonization are few: we have only come across a single study reporting AMF responses to clipping and water manipulation treatments under field conditions (Allen et al. 1989). The study used two species of grasses with differing herbivory tolerance, subjected to once-a-year clipping treatments over four years, and soil water availability manipulations (using rain-out shelters and irrigation treatments) over two years. The study found no consistent patterns of AMF colonization responses to either treatment, and was thus unable to provide definite answers to the question of top-down versus bottom-up controls on plant-AMF interactions.

Here, we report results from experiments conducted in Trans-Himalayan rangelands in Spiti, Himachal Pradesh, India, a high-altitude, cold and semiarid grazing ecosystem. We tested the predictions regarding the influence of top-down and bottom-up forces on AMF colonization (Fig. 1) by manipulating soil water availability and aboveground biomass loss. Irrigation experiments have shown that plants in this region are primarily water limited, with plant production increasing with water supplementation in the absence of herbivory (Bagchi & Ritchie 2011). Hence, we used water to test the predictions of bottom-up forces in this study. Specifically, we predicted that (i) conditions of high water availability (such as under short-term water supplementation, or years with relatively high annual precipitation) will be associated with low AMF levels, and (ii) AMF colonization levels will not be responsive to aboveground tissue loss (either due to short-term chronic clipping or long-term vertebrate grazing), given the greater influence we predict bottom-up forces to have on AMF colonization. We tested the predictions using four graminoids species for the experiments. While the host species can be expected to differ quantitatively in their responses to changes in water availability and aboveground tissue loss, we still expected to see qualitative similarity between them so that bottom-up forces influence AMF colonization levels more strongly than top-down forces.

## Methods

### Study area

The study was carried out in the rangelands around Kibber village (32° N 78° E) in the Spiti region, a 12,000 km2 area that is part of the arid to semiarid Trans-Himalayan landscape. The area receives low precipitation; rainfall between 2012 and 2014 averaged ~36 mm, and snowfall in the three years averaged ~340 cm (as recorded at the nearest weather station in Kaza, ~12 km away). During this study, years with high rainfall recorded lower snowfall and vice versa (Fig. 2).

**Fig. 2.**
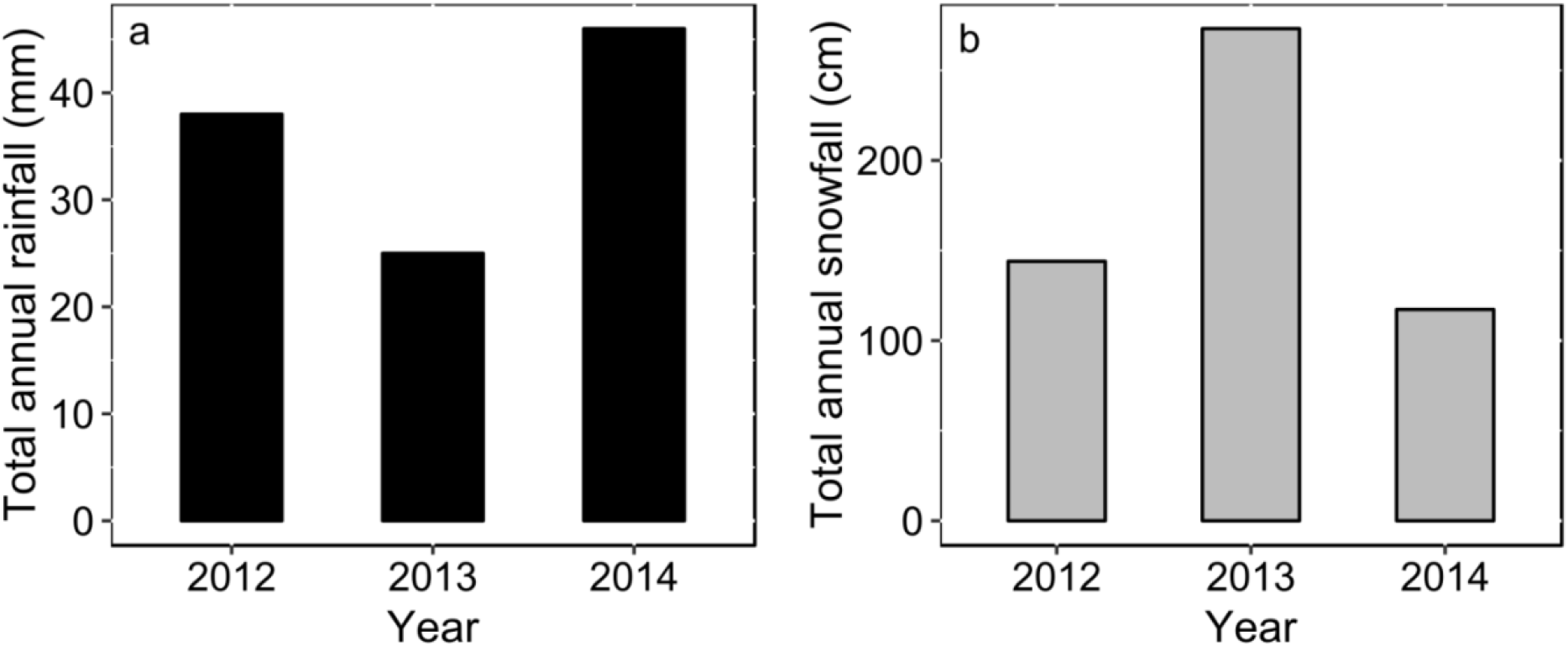
Total annual rainfall and snowfall from 2012 to 2014 in the Spiti region in the Trans-Himalaya, Himachal Pradesh, India.

The vegetation comprises perennial grasses (such as *Stipa* spp*., Elymus* spp.*, Leymus secalinus, Festuca* spp. and *Poa* spp.), sedges (such as *Carex* spp. and *Kobresia* spp.), herbs and a few shrubs (the dominant shrub being *Caragana versicolor*); the tree layer is absent. The growing season is short, from early May to end August, reflecting temperature and soil moisture patterns in the region (Bagchi et al. 2017). Further, vegetation is limited due to low water availability coupled with high altitude and cold temperatures, with a net annual aboveground plant production of 47.8 ± 2.2 g m-2 (Bagchi & Ritchie 2010b). However, these rangelands support a sizeable population of herbivores, both wild (such as bharal, *Pseudois nayaur,* and ibex, *Capra sibirica*) and domesticated (such as cattle, yak, goats, sheep, donkeys etc.). Wild herbivores and livestock consume ~60% of the plant production during the growing season, in watersheds primarily used by either herbivore type (Bagchi & Ritchie 2010a; Bagchi et al. 2012). A more detailed account of the study area is available in Bagchi et al. (2012).

### Experiment 1: Response of mycorrhizal colonization to clipping and supplemental water addition

We conducted a factorial clipping and watering experiment in July and August 2013 to assess the combined effect of simulated grazing and increased water availability on AMF colonization levels. We selected three species of grasses (*Elymus longae-aristatus*, *Leymus secalinus* and *Stipa orientalis*) and a sedge (*Carex melanantha*) as the focal host species for AMF, given their widespread nature and their role as common forage species for herbivores in the region (Mishra et al. 2004). We conducted the experiment within herbivore exclosures, which are 10 m × 10 m chain link fences that exclude wild herbivores and livestock, set up in 2005-06 in different watersheds in the rangelands around Kibber (Bagchi & Ritchie 2010b). Each host species was sampled, based on occurrence, from a subset of 16 herbivore exclosures selected from four watersheds (4-10 exclosures per host species; median = 8). In each exclosure, 20 individuals of each species were tagged and randomly assigned to four treatments (clipping, watering, clipping and watering or control (no clipping or watering)), with five individuals per treatment. Individual tagged plants of the four focal host species in the clipping treatment were cut to half their height at weekly intervals for four weeks to simulate chronic loss of aboveground tissue. Plants in the irrigated treatment were provided with ~250 ml stream water in a 10 × 10 cm area demarcated around them twice a week for four weeks with 3-4 day intervals between each watering event. Plants were watered nine times over the course of the month, with each plant receiving a total of ~2250 ml of water (equivalent to ~225 mm rainfall; ~9 times the total rainfall recorded in the region in 2013, which was 25 mm), in order to remove any potential water limitation. Plant roots were harvested at the end of the experiment, stored in FAA (10% formaldehyde, 5% acetic acid, 50% ethanol) and transported to National Centre for Biological Sciences, Bangalore, India, for AMF colonization estimation (methods described below). Individual plants were uprooted and roots still attached to the plant were then collected. Excluding samples where we could not harvest enough roots for AMF estimation, and depending on host species availability, we had data from 16-39 individuals (median = 26.5) per species per treatment, for a total of 427 samples.

Mycorrhizal responses, in terms of amount of root colonization, to the factorial clipping and watering treatments as compared to the control treatment (with neither clipping nor watering) in the four focal host species were analysed using generalized linear mixed models (GLMM) with binomial errors, given that AMF colonization data were in the form of proportions, and exclosure identity was used as the random factor to account for inter-exclosure variation in data (Bolker et al. 2009). Individual plants were the sampling unit, and treatment, host species and their interactions were used as predictors, with AMF colonization as the response variable. Differences between treatments and control allowed us to test predictions about the relative strengths of top-down and bottom-up forces in influencing AMF colonization levels, with the interaction term indicating whether different host species differed in AMF colonization responses to the different treatments.

### Experiment 2: Mycorrhizal colonization responses to long-term herbivory exclusion

This experiment tested the prediction that long-term grazer exclusion will not have changed AMF colonization levels significantly compared to plants in the adjacent grazed areas. This follows from the prediction that in resource limited conditions, like our water-limited study system, AMF colonization in plant roots will be high irrespective of grazer pressure (Fig. 1, Regions I and II). In August 2012 and August 2014, we collected individuals of each of the four host species from 8-12 (median = 9) and 5-6 (median = 5.5) exclosures and adjacent grazed (control) plots respectively, depending on the availability of each species. From each exclosure and control plot, roots from 5-10 individuals of each of the four host species were collected and stored in FAA for subsequent AMF colonization estimation. Excluding samples without enough roots for AMF estimation, we had data from 27-85 individuals (median = 56.5) per species per treatment in 2012 for a total of 447 samples, and 21-42 individuals (median = 30.5) per species per treatment in 2014 for a total of 248 samples.

Like for Experiment 1, mycorrhizal colonization responses to grazing as compared to ungrazed conditions in the four focal host species were analysed using GLMMs (Bolker et al. 2009). Individual plants were the sampling unit, and treatment (grazed/ungrazed), host species and their interactions were used as predictors, with AMF colonization as the response variable; exclosure identity was used as the random factor. Comparing AMF colonization levels in grazed and ungrazed plots across species in both years allows us to test the prediction that top-down forces have limited influence on AMF colonization levels.

### Mycorrhizal colonization responses to changes in annual precipitation levels

To test the prediction that AMF colonization varies negatively with annual precipitation, we used AMF colonization data collected over three years (2012-2014) and looked at how they varied depending on annual rainfall/snowfall for those years. Average AMF colonization per species in each of the three years was calculated using data from individuals within the exclosures in 2012 and 2014, and from the control treatment in the factorial experiment in 2013, since the experiment in 2013 was also done within the exclosures. Total annual rainfall and snowfall were quantified from September of the previous year to August of a given year to capture growing season (early May to end August) water availability at our study site.

We again used GLMMs for these analyses, with total annual rainfall and snowfall as predictors in two separate models, along with host species and their interaction with total annual rainfall/snowfall. Average AMF colonization of each species per year was used as the response variable in both models and exclosure identity was the random factor. Rainfall and snowfall values were log transformed before the analyses to meet normality assumptions. The relationship between rainfall/snowfall and AMF colonization levels across species tests the prediction that AMF colonization levels are strongly influenced by bottom-up forces.

The R package *lme4* was used to conduct all the GLMMs, while *lmerTest* was used to perform t-tests using Satterthwaite approximations on the degrees of freedom and *car* to perform Wald chi-square tests to assess the statistical significance of the fixed effects (Bates 2010; Bates et al. 2014, Kuznetsova et al. 2015; Bates et al. 2017). All analyses and figure preparation were carried out using R, version 3.2.4 (The R Foundation for Statistical Computing Platform, 2016).

### Arbuscular mycorrhizal fungal colonization estimation

Roots for AMF estimation from these experiments were collected by uprooting individual plants, ensuring that only roots still attached to the plant were collected after uprooting. This consideration also limited the amount of roots that could be harvested per individual, and thereby the amount of root available for AMF colonization estimation. The depth to which the soil was dug varied with species and individuals. To estimate AMF colonization in individual plants, we treated 1-2 cm pieces of the fine roots with 10% KOH at 90 °C for 1 hr and then stained them using the ink and vinegar method (Vierheilig et al. 1998, 2005): KOH treated roots were washed, treated with a 5% solution of black ink (Parker Quink, Bangalore, India) in 5% acetic acid at 90 °C for 15 min, and destained with distilled water acidified with a few drops of acetic acid. AMF colonization was estimated using the grid line intersection method (Giovannetti & Mosse 1980). Briefly, intersections of the stained fine roots with grid lines on a plate were observed under 400× magnification and scored for presence of arbuscules, vesicles and/or aseptate hyphae. AMF colonization was calculated as the ratio of the number of intersections that showed AMF presence to the total number of intersections observed. Data from 11-12 intersections for each sample were used to calculate AMF colonization, following estimation of the optimum effort in terms of number of total intersections observed required for arriving at accurate AMF colonization measures (details in Appendix I). AMF colonization measurements from individuals with data from 33-45 (median = 41.5) intersections showed that colonization estimates stabilized at ~10 intersections.

## Results

Clipping and watering treatments elicited different AMF colonization responses in each species. In the long-term grazer exclusion experiment too, AMF colonization responses differed with host species and across years. Annual rainfall/snowfall, however, influenced AMF colonization levels similarly across host species (Table 1).

**Table 1.** Summary of statistical analyses assessing AMF colonization responses to short-term clipping and watering treatments in the factorial experiment, long-term grazer exclusion and to total annual precipitation levels. *P*-values below 0.05 are in bold.

| Experiment | Effect | Wald chi-square | df | *P* |
| --- | --- | --- | --- | --- |
| Factorial clipping and watering | Treatment | 6.406 | 3 | 0.093 |
|  | Host species | 119.550 | 3 | < **0.001** |
|  | Interaction | 12.236 | 9 | 0.20 |
| Long-term herbivory exclusion (2012) | Treatment | 0.8336 | 1 | 0.36 |
|  | Host species | 29.0801 | 3 | < **0.001** |
|  | Interaction | 35.8726 | 3 | < **0.001** |
| Long-term herbivory exclusion (2014) | Treatment | 0.8339 | 1 | 0.36 |
|  | Host species | 189.1219 | 3 | < **0.001** |
|  | Interaction | 21.6219 | 3 | < **0.001** |
| Annual precipitation effects: |  |  |  |  |
| (i) Total annual rainfall | Rainfall | 344.242 | 1 | < **0.001** |
|  | Host species | 38.073 | 3 | < **0.001** |
|  | Interaction | 19.125 | 3 | < **0.001** |
| (ii) Total annual snowfall | Snowfall | 283.526 | 1 | < **0.001** |
|  | Host species | 38.858 | 3 | < **0.001** |
|  | Interaction | 17.630 | 3 | < **0.001** |

The effects of experimental clipping and watering treatments on AMF colonization differed between host species. In *Carex*, AMF colonization decreased by ~11.8% in the watered + clipped treatment (*P* = 0.032), and in *Stipa*, colonization levels increased in the watered treatment by ~5.9% (*P* = 0.034), compared to levels of AMF colonization in the control treatment in either species. None of the treatments had any effect on AMF colonization for the other host species (Fig. 3).

**Fig. 3.**
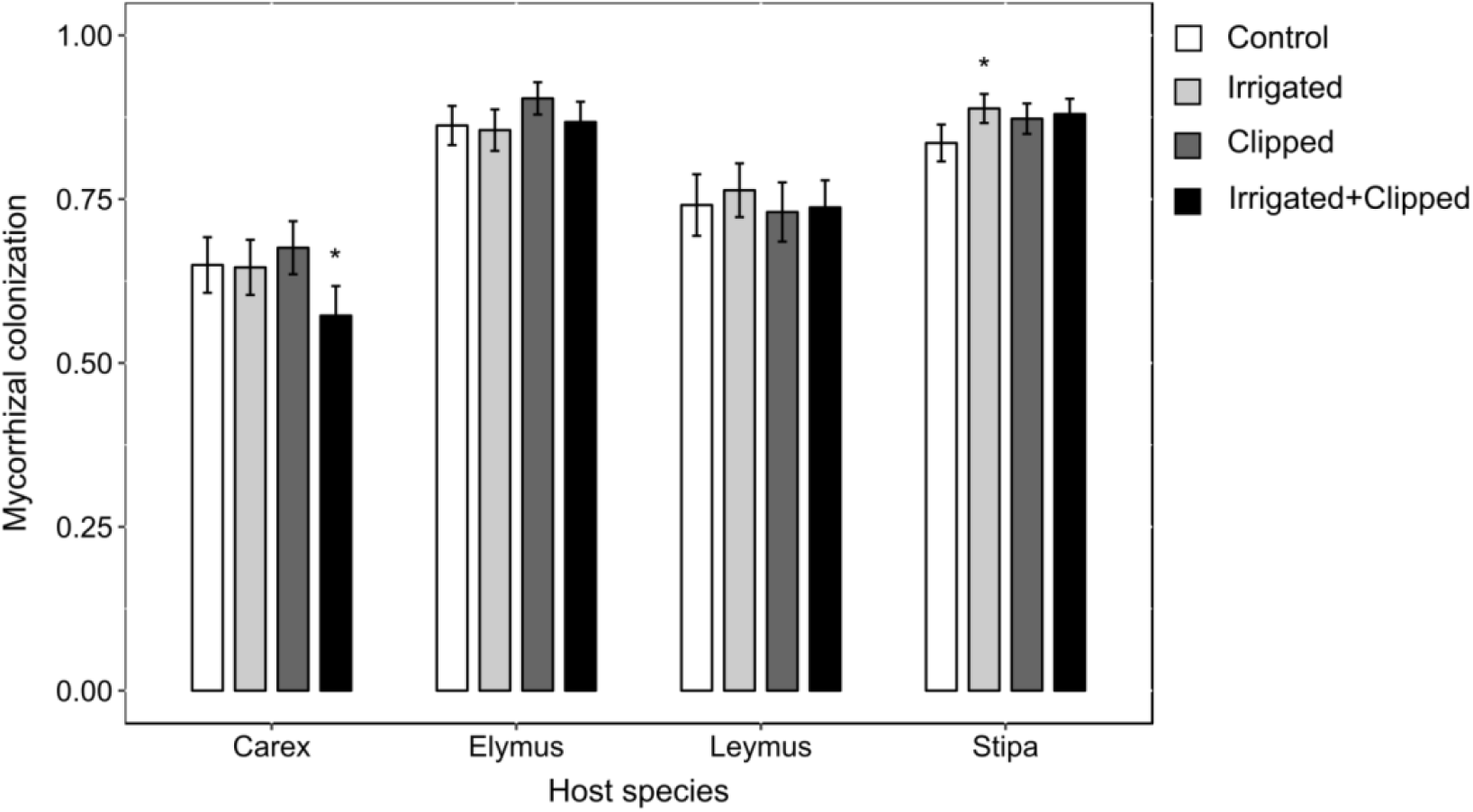
AMF colonization responses to short-term clipping and watering. Bars are species averages and error bars represent 1 SE, obtained from fixed effects statistics in the mixed effects model used for analyses. Asterisks indicate mixed effects model output of statistically significant within species differences between clipping and watering treatments and the control (* *P* < 0.05).

Long-term grazer exclusion effects on AMF colonization levels differed among host species. In 2012, AMF colonization levels in the grazed plots were lower by ~4.5% (*P* = 0.01) and ~10.2% (*P* < 0.001) when compared to the ungrazed plots in *Carex* and *Elymus* respectively, but were higher in the grazed plots by ~6.7% (*P* < 0.01) in *Stipa*. AMF colonization levels were unaffected by grazing in *Leymus* (Fig. 4a). However, in 2014, a year with greater total annual rainfall (Fig. 2), AMF colonization levels in *Elymus* were higher by ~21.6% (*P* = 0.036), but lower by ~25.8% (*P* < 0.001) in *Stipa* in grazed plots compared to ungrazed plots, while the other host species showed no difference in AMF colonization levels between the two treatments (Fig. 4b).

**Fig. 4.**
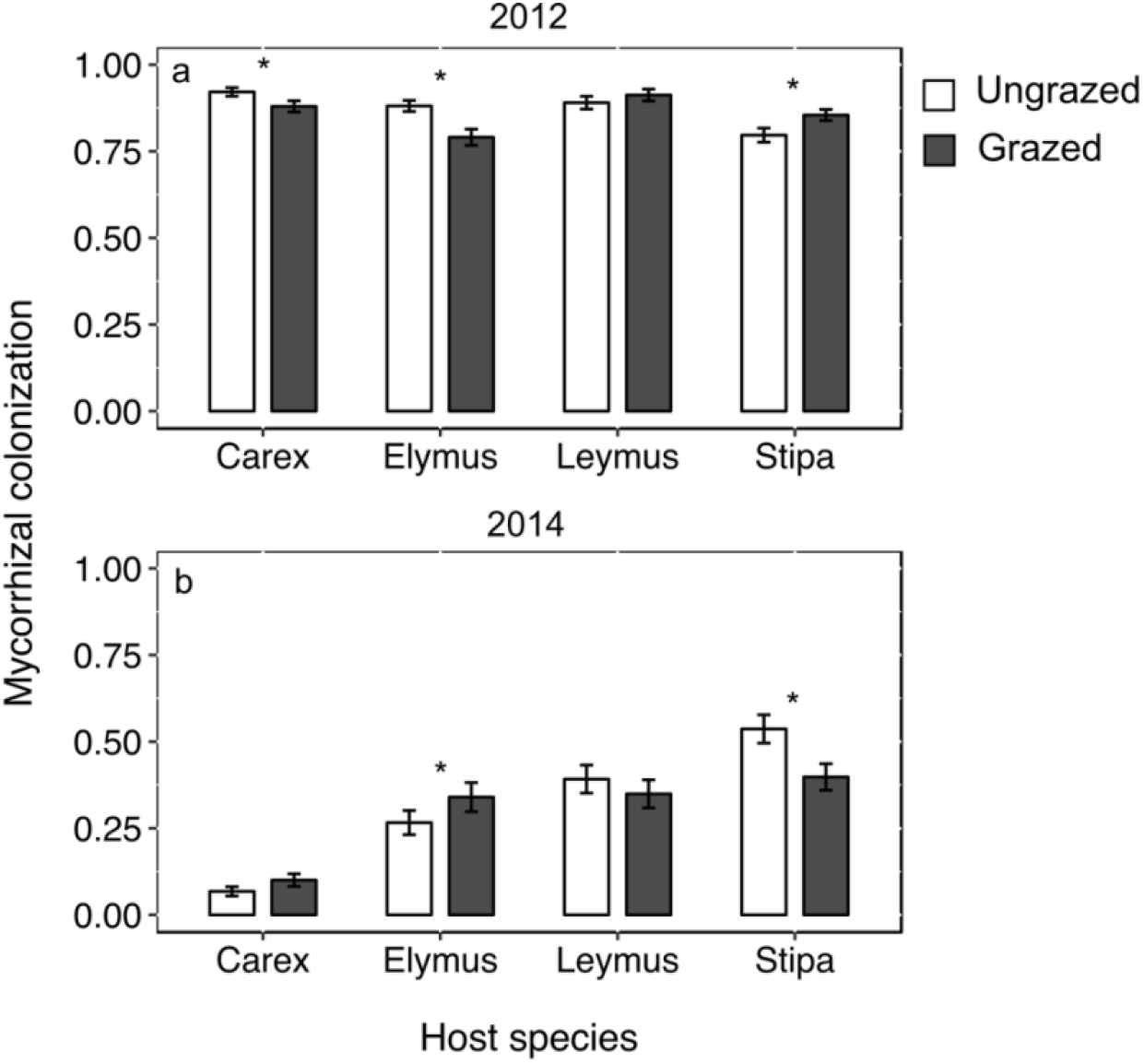
Long-term grazer exclusion experiment: chronic vertebrate grazing effects on AMF colonization levels in four host species in 2012 (a) and 2014 (b). Bars are species averages and error bars represent 1 SE, obtained from the respective fixed effects statistics in the mixed effects models used for analyses. Asterisks indicate mixed effects model output of statistically significant within species differences between treatments in each year (* *P* < 0.05, ** *P* ≤ 0.01, *** *P* < 0.001).

AMF colonization in all four host species varied negatively with total annual rainfall (*P* < 0.001) (Fig. 5a). However, since years with high total annual rainfall were characterized by low total annual snowfall and vice versa during the study (Fig. 2), AMF colonization was positively related to total annual snowfall (*P* < 0.001) (Fig. 5b).

**Fig. 5.**
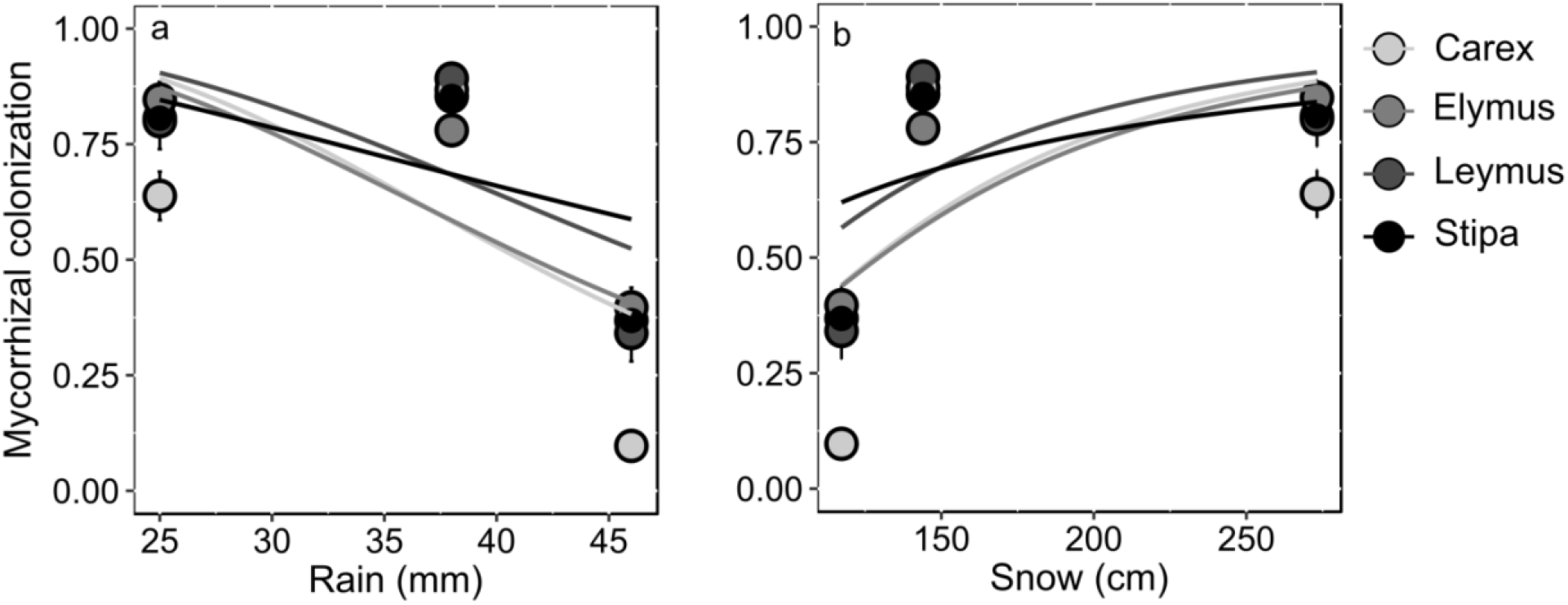
AMF colonization levels across years are related to total annual rainfall (a) and total annual snowfall (b). Lines are species wise slopes from the respective mixed effects models. Error bars represent 1 SE about the species means of AMF colonization levels.

## Discussion

Overall, one of the clearest findings from this study is that host species differ in AMF colonization responses to clipping and short-term changes in soil water availability. Long-term grazer exclusion effects on AMF colonization also differed between host species. AMF colonization levels also, however, differed across years. Given that water is the primary limiting factor for plant growth in this region, some of this variation in AMF levels across years, at least, must be explained by changing precipitation levels. AMF colonization decreased with annual rainfall lending support the prediction in Fig. 1, though with total annual snowfall we observed the opposite pattern. However, the fact that annual rainfall/snowfall effects were uniform across host species points to a potential stronger effect of resource availability in these semi-arid rangelands (see Appendix II). Further longer term studies are needed to gain a better understanding of the relative strengths of grazer and plant soil resource availability effects on AMF colonization levels.

Exclusion of chronic grazing by vertebrate herbivores had variable effects on AMF colonization in this study, differing with host species and across years. Variable effects of defoliation on AMF colonization have been widely reported in literature: ~53% of the studies in a recent review of grazing effects on mycorrhizae reported decreases in mycorrhizal colonization after defoliation, while the remaining reported increases or no changes (Barto & Rillig 2010). Experiments have found defoliation mediated reductions in AMF hyphal colonization in perennial grasses, especially grazing intolerant species such as *Themeda* (Allsopp 1998), and cucurbits (Barber et al. 2012), or increases as in an experiment with a species of bunchgrass (Wallace 1987). Results from heavily grazed systems such as the Serengeti have also shown that long-term herbivory exclusion effects on AMF are very variable (Antoninka et al. 2015), while other studies on perennial grasses conducted under field conditions often show no effect of defoliation on mycorrhizal colonization (Barto & Rillig 2010).

AMF colonization responses to chronic grazing by livestock and wild herbivores were different from responses to short-term clipping in this study. This may be because the level (or duration) of clipping we employed may not have been sufficient, or because clipping does not exactly mimic herbivory and C loss due to defoliation alone does not drive AMF colonization changes. Studies on species of forage grasses have shown that while heavy defoliation might decrease AMF colonization levels, there is no effect at intermediate levels (Bethlenfalvay et al. 1985). Further, in a study in a tallgrass prairie ecosystem, mowing did not elicit changes in AMF colonization levels (Eom et al. 1999), whereas increased AMF colonization was reported under cattle grazing (Eom et al. 2001).

This study found that AMF colonization varied with annual precipitation levels, supporting the prediction that resource availability is a potential predictor of AMF colonization levels. Precipitation and AMF colonization data from more years are needed to confirm whether AMF colonization levels in this region consistently respond to precipitation levels. However, other studies in semi-arid and mountain grasslands have also found inter-annual precipitation differences or seasonal changes in soil moisture to be stronger drivers of plant performance (Giese et al. 2011), as well as AMF colonization responses (Lugo et al. 2003) than grazing. More generally, several studies involving defoliation and resource manipulation support the prediction of a stronger effect of resource availability than herbivory on AMF colonization. A recent study conducted in the Serengeti found no evidence for grazing effects on several AMF metrics such as hyphal standing crop in the soil, new hyphal production and spore abundance, while rainfall and soil parameters such as P and organic matter content were found to be important predictors of AMF (Antoninka et al. 2015). Studies have indicated that even herbivore mediated effects on AMF colonization might in fact be due to herbivore effects on soil parameters (Ba et al. 2012). However, AMF colonization responses to altered resource availability can be extremely variable (Mohan et al. 2014). Indeed, studies have reported increases or decreases in AMF levels under drought (as reviewed in Mohan et al. 2014) or no changes (e.g. Wallace 1987; Busso et al. 2008) in AMF colonization levels, suggesting that factors other than resource availability may also play a role in influencing AMF levels in plant roots.

AMF colonization levels and their responses to top-down and bottom-up forces differed between host species in this study. Indeed, some earlier studies have also reported host species to explain more of the variation in AMF colonization levels than defoliation treatments (Walling & Zabinski 2006). AMF colonization levels have been demonstrated to be variable even between species belonging to the same genus (Busso et al. 2008). Host species variation in mycorrhizal responses can be attributed to possible differences in mycotrophy (Gange et al. 2002), nutrient demand (Fosaa & Olsen 2007), or differences in competitive ability in general (Busso et al. 2008), or grazing induced changes in the AMF community itself (Ba et al. 2012; Antoninka et al. 2015).

Several studies involving defoliation, fertilization or resource gradients have found a stronger influence of bottom-up rather than top-down forces on AMF colonization levels (e.g., Eom et al. 1999; Antoninka et al. 2015), supporting the predictions in Fig. 1. However, studies that test the relative strengths of these factors, especially when the limiting resource for plant growth is water, are scarce. In this study, we find that in arid high-altitude rangelands precipitation levels potentially influence AMF colonization levels. However, the effect of water availability and herbivory differ with host species, making it challenging to predict the effects of changing grazing and precipitation regimes on plant-AMF interactions. Further, long-term herbivore exclusion experiments in the Spiti region have shown that grazing alters vegetation production and composition (Bagchi & Ritchie 2010a), leads to more fungus-dominated soil communities, and reduces microbial biomass and potential microbial respiration in the region (Bagchi et al. 2017). However, whether these changes in vegetation composition, production and soil factors influence community level AMF colonization in plant roots is not yet known. Long-term grazer exclusion/clipping and water manipulation experiments that also take in account AMF and vegetation community shifts are essential to get a comprehensive picture of controls on plant investment in AMF, and consequently C and nutrient storage and cycling, in these Trans-Himalayan systems.

## Acknowledgements

We are grateful to the Himachal Pradesh Forest Department for permission to conduct field work, especially to the Principal Chief Conservator of Forests, the Divisional Forest Officer, the Range Forest Officer, and Mr. Satpal Dhiman for their continued support. We thank Dorje Chhering, Dorje Chhewang, Tanzin Chhewang and Kunga Thargye for help with field work, and Rinchen Tobge, Tanzin Thinley and others at Kibber for their hospitality and friendship.

We also thank Shishira Suresh and Tanvi Somiah who helped with a part of the lab work. National Centre for Biological Sciences, Bangalore, provided core funding for this study.

## Appendix I Estimating optimum effort for arbuscular mycorrhizal fungal colonization measurement

Arbuscular mycorrhizal fungal (AMF) colonization levels were measured using the grid line intersection method, where intersections of the stained fine roots with grid lines on a plate were observed under 400× magnification and scored for presence of arbuscules, vesicles and/or aseptate hyphae. AMF colonization was calculated as the ratio of the number of intersections that showed AMF presence in the root samples observed to the total number of intersections observed.

In this study, the median number of root pieces observed per sample ranged from 8 to 16 (overall median = 11.5). The total intersections observed per species per year had medians ranging from 11 to 32 (overall median = 16.5). We used data from 11-12 intersections for each sample to calculate AMF colonization, since only limited amount of roots could be harvest in the field while still making sure it belonged to an individual plant.

Optimum effort to arrive at accurate AMF colonization measures was estimated using individuals of the four focal host species in this study from which we could harvest the most roots, and thereby obtain AMF presence/absence data from 33-45 (median = 41.5) intersections. Data for total intersections observed (and the corresponding data for the number of intersections with AMF infection) were sampled without replacement over 100 iterations, with increasing number of intersections observed (from ~1 intersection to the maximum number of intersections available per sample), and the AMF colonization level was calculated in each case. In all cases, the simulations revealed that AMF colonization estimates stabilize at ~10 intersections (Fig. 1).

**Fig. 1.**
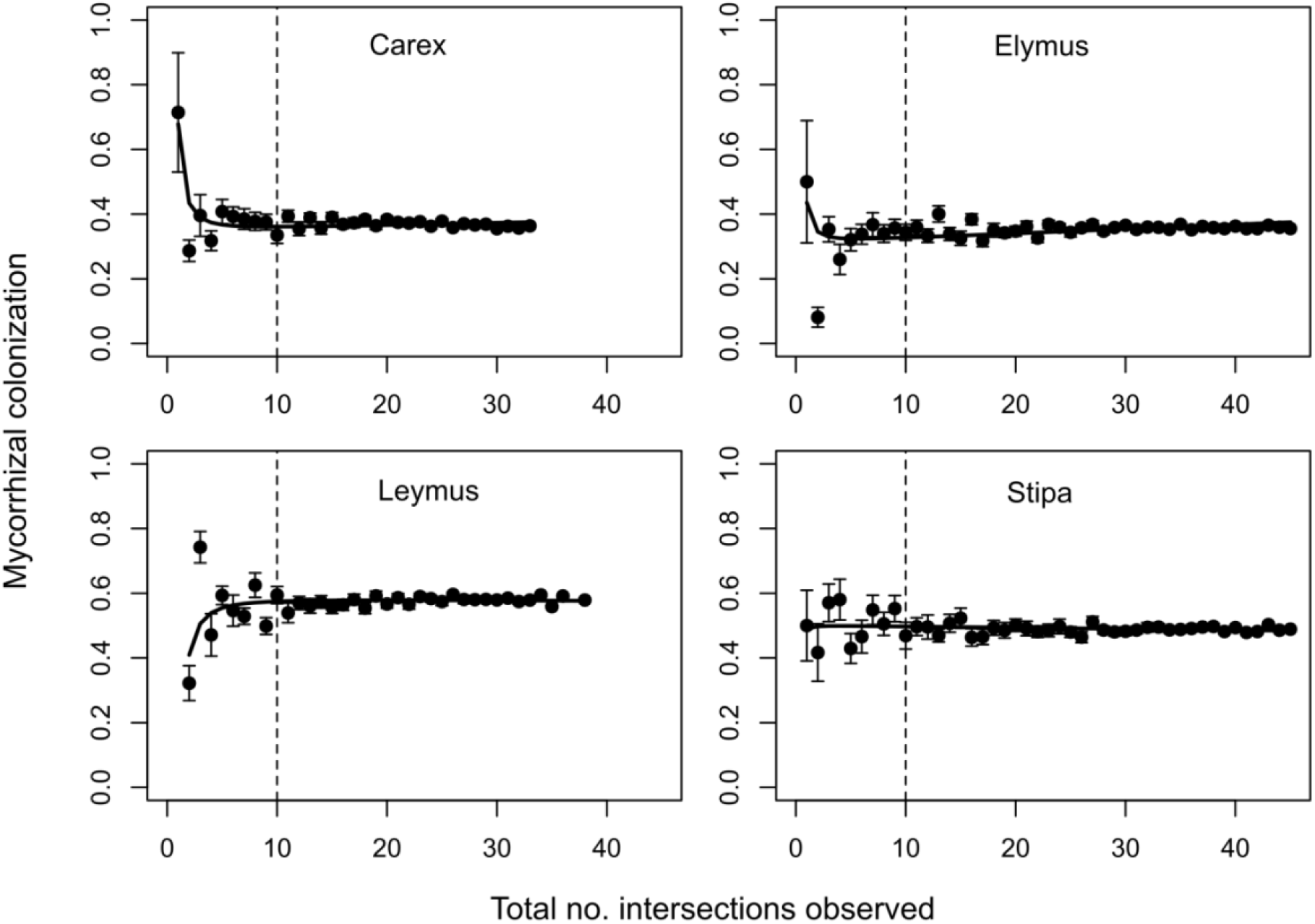
Relationship between total numbers of intersections observed and AMF colonization estimation for individuals of the four focal host species used in this study. The points represent AMF colonization estimated from 1 to the maximum number of intersections data was available for, averaged over several iterations. Error bars are 1 SE about the mean. The dotted line represents 10 total intersections observed. We used 11-12 intersections per sample to calculate AMF colonization levels in this study.

## Appendix II Meta-analytical assessment of AMF responses to short- and long-term changes in grazer pressure and soil water availability

In this study, we studied top-down vs. bottom-up controls on AMF colonization levels using short-term clipping and watering experiments, long-term grazer exclusion, and colonization responses to annual rainfall and snowfall over a three-year period. Here we present results from a meta-analysis, where we compare AMF colonization responses to each of these experiments, in order to capture the overall learnings from this study.

## Methods

The methods for this meta-analysis follow the procedure described in Hedges et al. (1999) and earlier meta-analyses of AMF responses to various drivers (such as Treseder 2004).

There are six ‘experiments’ in this study, looking at aboveground biomass loss and/or soil water availability effects on AMF colonization (Table 1): (a) Long-term grazing (for which individuals within long-term grazer exclosures served as controls), (b) short-term clipping, (c) short-term irrigation, (d) short-term clipping combined with irrigation (for b - d, all plants were inside long-term exclosures, and plants that were neither clipped nor watered served as controls), (e) annual rainfall from 2012 to 2014, and (f) annual snowfall from 2012 to 2014 (for e and f, years with the lowest rainfall (2013) and lowest snowfall (2014), served as the controls, respectively, while the other two years were considered the ‘experiment’).

For the long-term grazing experiment, the four focal species, individual exclosures and the two years the data were collected in were considered as separate ‘studies’ in the meta-analysis, for a total of 48 studies. In other words, data from one host species, in one exclosure for a given year made up one ‘study’. Similarly, for the short-term clipping and irrigation experiments (experiments b - d above), data from each focal species and individual exclosure were considered as separate studies, for a total of 21, 24 and 26 studies respectively (the numbers vary because we used only data from those species and exclosures that could be matched to data from the control group, and due to sample loss and other reasons described in the main text, not all exclosures had data for all treatments). For the annual rainfall and snowfall experiments (experiments e and f above), data from each species and year were considered as separate studies (for a total of 8 ‘studies’ each). Further division of studies by exclosure identity was not possible for the rainfall and snowfall data because of violation of the assumption of normal approximation to the sampling distribution of the log response ratios (calculation of which is described below). Normal approximation was checked by making sure that the majority of the studies had 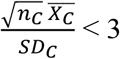, where *n*_*c*_ is the number of replicates in each study in the control group, 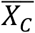 is the mean AMF colonization, and *SD*_*c*_ is the standard deviation, and similarly for studies in the experiment group (Hedges et al. 1999).

For each of the ‘studies’, effect sizes were calculated as the natural log of the response ratio (R), which is the ratio of average AMF colonization levels in the experiment over control. Further, within-study variance (*v*_ln R_), which is a function of mean, standard deviation and number of replicates in the experiment and control groups were estimated. These, along with estimates of the between-study variance, were then used to calculate the weighted mean of the log response ratios, and the CIs associated with the weighted mean, for each experiment. Here, we present the response ratio calculated from the weighted mean log response ratio, associated confidence intervals (CI), the Q statistic which captures the heterogeneity among studies, and associated *P*-values (Hedges et al. 1999).

## Results

Short- and long-term grazer pressure have no effect on AMF colonization levels in the semi-arid rangelands that we studied (Fig. 1). Short-term increases in water availability, likewise, does not have an effect. However, increased annual rainfall is associated with decreases in AMF colonization levels in the focal host species. In the three years the study spanned, years with high rainfall were associated with low snowfall and vice-versa. Increased snowfall is associated with increased AMF colonization in graminoids in this region (Fig. 1). Further, AMF colonization responses appear very sensitive to host species identity, and responses also varied from year to year (see Results section in the main text of Chapter 2). This is also reflected in the Q statistic values for all the experiments, which shows that there is significant variation between studies in all cases (Table 1).

**Fig. 1.**
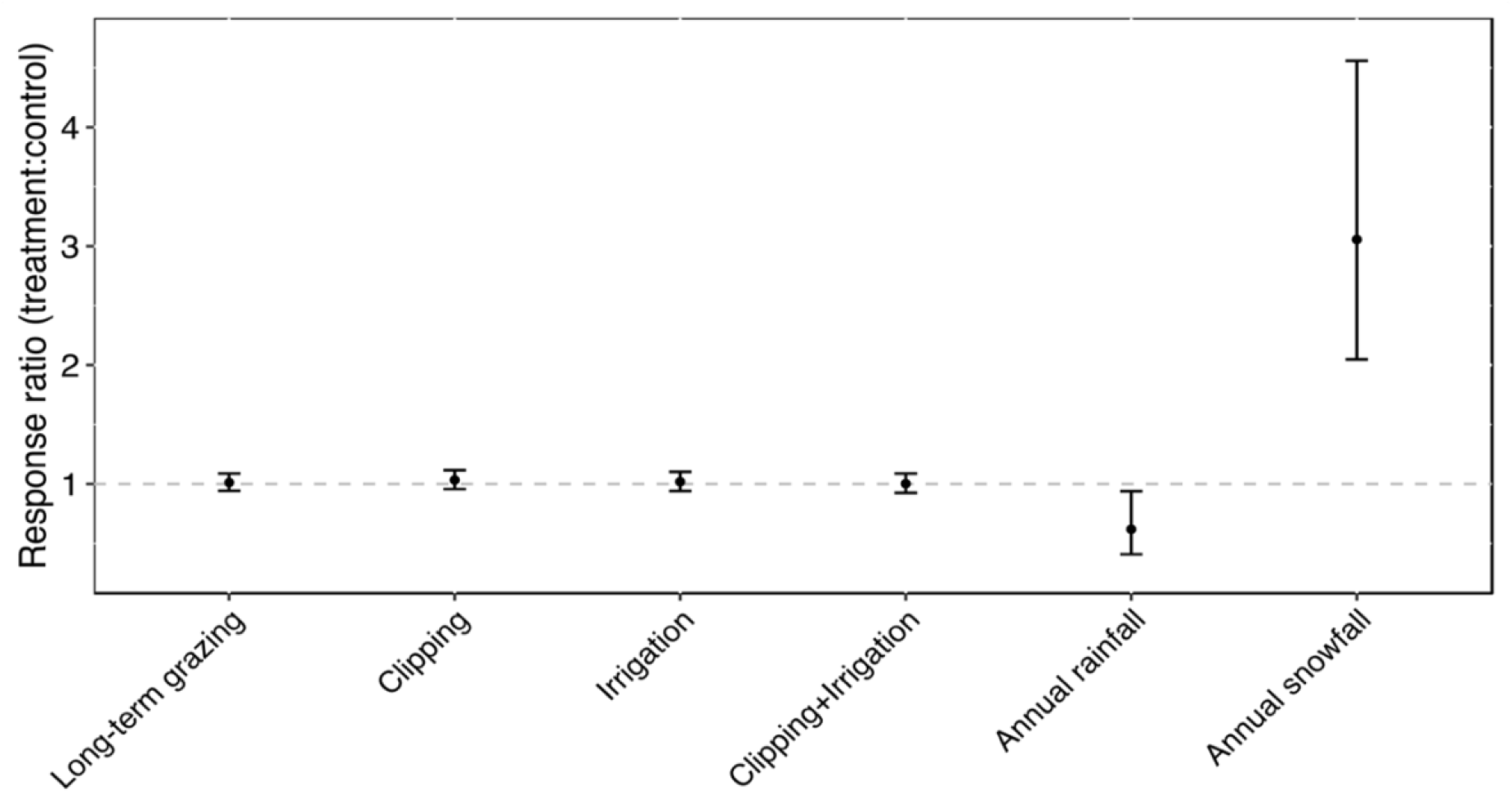
Weighted mean response ratios of AMF colonization responses to long- and short-term aboveground biomass loss, short-term irrigation and annual rainfall and snowfall. The grey dashed line represents a response ratio of 1, or no difference between AMF responses between experiment and control groups. Error bars are CIs.

**Table 1.** Weighted mean response ratios, associated CIs, Q statistics, degrees of freedom (df) and *P*-values for AMF colonization responses to long- and short-term aboveground biomass loss, short-term irrigation and annual rainfall and snowfall.

| Experiment | Control | Response ratio (CI) | Q | df | P |
| --- | --- | --- | --- | --- | --- |
| Long-term grazing | Long-term grazer exclusion | 1.01 (0.94, 1.09) | 242.80 | 47 | <0.001 |
| Clipping | No clipping or irrigation | 1.03 (0.96, 1.11) | 37.98 | 20 | 0.009 |
| Irrigation | No clipping or irrigation | 1.02 (0.94, 1.10) | 40.36 | 23 | 0.014 |
| Clipping + irrigation | No clipping or irrigation | 1.00 (0.92, 1.09) | 47.10 | 25 | 0.005 |
| Annual rainfall | Lowest rainfall year (2013) | 0.62 (0.41, 0.94) | 501.21 | 7 | <0.001 |
| Annual snowfall | Lowest snowfall year (2014) | 3.05 (2.05, 4.56) | 306.30 | 7 | <0.001 |

